# Inhibition of neutral sphingomyelinase-2 facilitates remyelination

**DOI:** 10.1101/686287

**Authors:** Seung-Wan Yoo, Amit Agarwal, Matthew D. Smith, Saja S. Khuder, Emily G. Baxi, Ajit G. Thomas, Camilo Rojas, Mohammed Moniruzzman, Barbara S. Slusher, Dwight E. Bergles, Peter A. Calabresi, Norman J. Haughey

## Abstract

For reasons that are not completely understood, remyelination is often incomplete, producing thin myelin sheaths with disorganized structure. We investigated the cellular basis for this altered myelin structure, and found that the response of oligodendrocyte progenitor cells (OPCs), and mature oligodendrocytes to TNFα and IL-1β is modified by the expression of the sphingomyelin hydrolase nSMase2. OPCs do not express nSMase2, and exhibit a protective response to these cytokines manifest by decreased ceramide, increased sphingosine 1-phosphate, and increased cell motility. Mature oligodendrocytes express nSMase2, and respond to TNFα and IL-1β with a stress phenotype, evidenced by increased ceramide, decreased sphingosine, and active caspase 3. Pharmacological inhibition or a targeted genetic deletion of nSMase2 *in vivo* increased myelin thickness, and enhanced myelin compaction. These results suggest that inhibition of nSMase2 improves the quality of new myelin by protecting maturing/myelinating oligodendrocytes. Pharmacological inhibition of nSMase2 following a demyelinating event could stabilize the structure of these newly formed myelin sheaths and protect them from secondary demyelination.

## Introduction

A highly synchronized and complex generation of multiple classes of lipid and select proteins regulates the biophysical properties of myelination. The lipid composition of myelin is asymmetrically distributed to allow for membrane curvature and myelin compaction. Intramembrane adhesions are critical for myelin to maintain its compact structure. The head to head arrangement of lipids in myelin creates a repulsive energy between opposing layers that is overcome at the major dense line by the stabilizing actions of myelin basic protein (MBP). Electrostatic interactions between negatively charged phosphatidyl serine (PS) head groups, and positively charged surface groups on MBP, combined with attractive van der Walls force creates a highly stable interface (Min, Kristiansen et al., 2009). Adhesion at the intraperiod line is less stable, and relies heavily on the interactions of proteolipid protein (PLP) with membrane lipids through van der Walls force to overcome repulsive energies originating from thermal undulations. Small changes in the lipid components can modify these adhesive forces and result in the decompaction and/or degeneration of myelin. We previously reported modifications in the lipid composition of normal appearing white matter (NAWM) at sights adjacent to lesions in multiple sclerosis (MS) brains, and calculated that this change in lipid composition would produce an order of magnitude increase in the repulsive energy between bilayers, that would destabilize myelin structure (Bandaru, Mielke et al., 2013). Although these results were compelling, using autopsy material we were unable to determine the mechanism(s) that contributed to these perturbations in the lipid content of regenerated myelin. In this study we show that pharmacological inhibition or genetic deletion of the sphingomyelin hydrolase neutral sphingomyelinase 2 (nSMase2) protects oligodendrocytes during re-myelination, resulting in a partial normalization of myelin lipid content and restoration of myelin compaction.

## Results

### Modifications in the lipid composition of myelin following remyelination

Compared with control mice fed a standard diet, we observed an 89 ± 9.5% reduction of black gold staining in the posterior corpus callosum (pCC) of mice fed a cuprizone containing diet for 4 weeks, consistent with a robust demyelination in this brain region. Four weeks after return to a standard diet, there was a considerable amount of remyelination in the pCC, but black gold staining remained 32 ± 4.3% below control levels (Fig 1A, B). Ultrastructural analysis of pCC showed similar results with a 51 ± 11.7% reduction in the number of myelinated axons following 4 weeks of cuprizone feeding, that partially recovered 4 weeks after return to standard diet.

**Figure 1.**
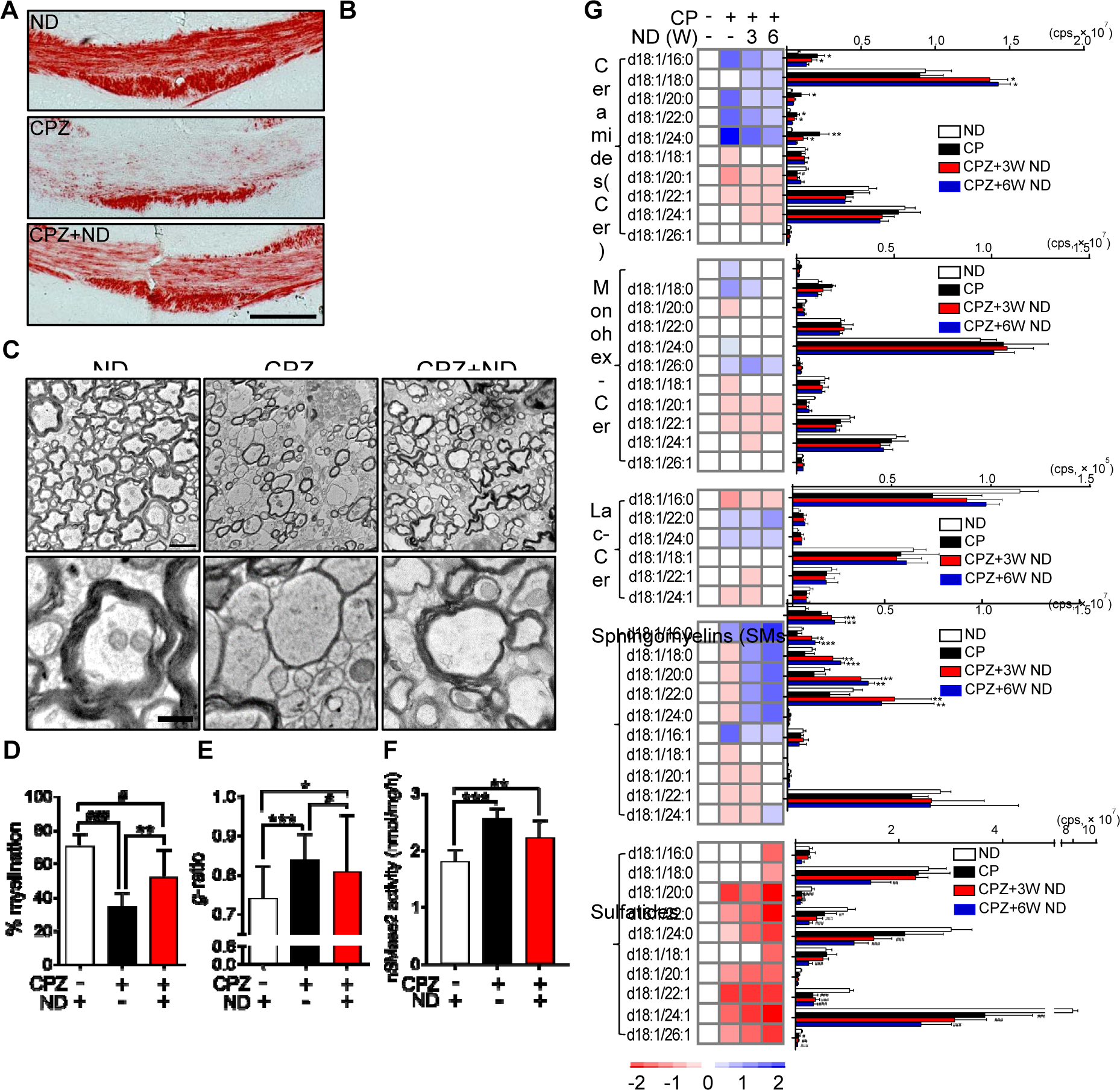
Modifications in the lipid content of remyelinated axons. A. Representative images of Black Gold staining in pCC of mice fed a normal diet (ND), mice fed a diet containing cuprizone (CPZ) for 4 weeks, and mice fed a CPZ containing diet for 4 weeks followed by a return to ND for 4 weeks (CPZ+ND). Images show extensive demyelination in the CPZ group, and a partial restoration of gross myelin structure in CPZ+ND group. Scale bar = 200 μm.
B. Quantitation of Black Gold staining in pCC of mice fed ND, CPZ, CPZ+ND (n=5 in each group).
C. Representative electron microscope images of the pCC in mice fed a ND, a CPZ diet, and a CPZ+ND diet. Images show extensive demyelination in the CPZ group, and partial restoration of myelin ultrastructure in the CPZ+ND group with many thinly myelinated axons with disorganized structure. Lower images are magnifications of the corresponding upper images. Scale bars = 2 μm in the top panel and 500 nm in the bottom panels.
D. Quantitation of the percent of myelinated axons in pCC after the indicated treatment conditions (n=100-250 axons from 3 animals per group).
E. Quantitation of g-ratios for myelin sheaths in pCC after the indicated treatment conditions (n=100-250 axons from 3 animals per group).
F. nSMase2 activity in pCC after the indicated treatment conditions (n=5 in each group).
G. Heat maps (left) and quantitative comparisons (right) of the indicated sphingolipids (ng/mg protein; n=5 in each group). Data show increases in multiple ceramides with decreases in sphingomyelins and sulfatides in the CPZ group, with time-dependent reductions of ceramides and increases is SMs in the CPZ+ND groups. Sulfatides do not recover in the CPZ+ND groups, and remain depleted.

Data information: Data are presented as mean ± S.D. * = *p* < 0.05, ** = *p* < 0.01, *** = *p* < 0.001, and # = p<0.05, ### = p<0.001 as indicated. ANOVA with Tukey post-hoc comparisons.

However, the number of myelinated axons remained 26 ± 23.0% below control (Fig 1C-E), and many axons appeared to be thinly myelinated compared with control mice (Fig 1C, E). Based on evidence from human brain tissue studies normal appearing white matter close to lesion sites contain a modified sphingolipid composition (Wheeler, Bandaru et al., 2008), we next measured how cuprizone modified sphingolipid content during demyelination, and at multiple time points during remyelination. After 4 weeks of a diet containing cuprizone we found that the pCC contained increased levels of multiple saturated ceramides (16:0, 20:0, 22:0, 24:0), reduced levels of multiple saturated sphingomyelins (C20:0, 22:0, 24:0), and a substantial reduction in multiple sulfatides (C20:0, 22:0, 22:1, 24:1, 26:1) (Fig. 1G). This pattern of change in lipid composition was similar (albeit to a lesser degree) in aCC, cortex, and hippocampus, and to a lesser extent in striatum and cerebellum (Supplementary Figure 1). After return to a standard diet, ceramide levels slowly returned to control levels over ∼6-weeks, with the exception of C18:0 ceramide (Fig 1G). The initial reduction of sphingomyelin after cuprizone feeding rebounded to levels increased above control, and multiple saturated sphingomyelins remained elevated 6 weeks after return to standard diet (Fig 1G). Decreased levels of sulfatides following cuprizone feeding did not recover within 6 weeks following return to a regular diet (Fig 1G). The pattern of increasing ceramides and decreasing sphingomyelins during demyelination is consistent with the action of nSMase2 that hydrolyzes sphingomyelin to ceramide. Indeed we found that nSMase2 activity increased by 52 ± 3.5% in pCC after 4 weeks of cuprizone feeding, and was reduced, but remained elevated 27 ± 16.5% above control levels 4 weeks after return to a regular diet (Fig 1F).

### Inflammatory cytokines protect OPCs, and promote apoptosis during OPC differentiation

Inflammation is functionally important for oligodendrocyte regeneration. Although the rate of OPCs proliferation in tissue culture was not modified by a 2-day treatment with TNFα or IL-1β (Fig 2A), a 4-day treatment of OPCs with TNFα or IL-1β reduced immunoreactivity for the active form of caspase3 (Fig 2B), consistent with a protective response to inflammatory cytokine challenge. TNFα also rapidly increased the rate of OPC migration (Fig 2C, Supplementary movie1, 2) suggesting that inflammatory challenge directly enhanced the migration of OPCs. We next exposed OPCs to TNFα and IL-1β every 2-days for 7-days during differentiation and found that the fraction of differentiated MBP+ oligodendrocytes was reduced from 42.5 ± 6.6% in control cultures (57 ± 32 cells/128 ± 58 cells) to 12.2 ± 1.8% with TNFα exposure (15 ± 3 cells/119 ± 2 cells), and to 15.9 ± 4.6% following IL-1β (17 ± 7 cells/106 ± 24 cells)(Fig 2D,E). We also found a dose related increase in the fraction of activated caspase3, with the highest dose of TNFα or IL-1β (100 ng/ml) producing a 324 ± 74.1%, and 318 ± 70.9% respectively increase in activated caspase3 staining (Fig 2F,G). These data suggest that inflammatory stimuli promote a protective/instructive response in OPCs that is modified to an injurious response as OPCs differentiate into MBP+ oligodendrocytes.

**Fig. 2.**
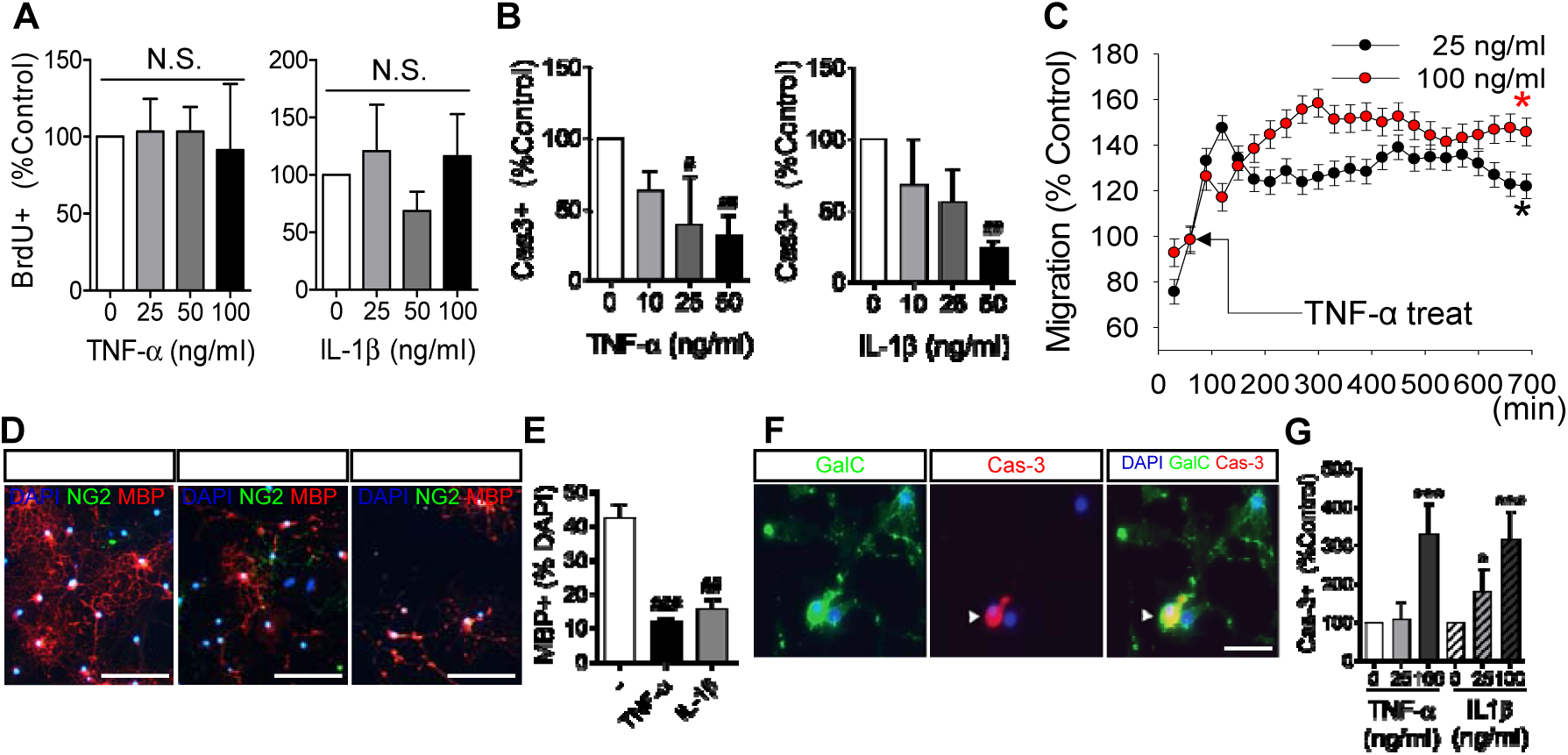
TNFα and IL-1β promote the survival and migration of oligodendrocyte progenitor cells, but stimulate cell death during the differentiation of progenitors to mature oligodendrocytes. A. Quantitation of oligodendrocyte progenitor cells (OPCs) proliferation by treatment with a dose response of TNFα (0-100 ng/ml), or IL-1β (0-100 ng/ml) as determined by incorporation of the uridine analog Bromodeoxy-Uridine (BrdU, 5 µg/ml) (n=3 in each group).
B. Quantitation of spontaneous cell death in OPCs by treatment with a dose response of TNFα (0-100 ng/ml), or IL-1β (0-100 ng/ml) for four consecutive days as determined number of active caspase-3 immunopositive cells (n=3 in each group).
C. Quantitation of OPCs migration over a 12h time frame showing increased cell migration following treatment with TNFα (25 or 100 ng/ml) (n=6 in each group).
D. Representative images of OPCs induced to differentiate by removal of PDGF. Cultures showed a reduction in the number of MBP+ cells after 7 days of treatment with TNFα or IL-1β (100 ng/ml) every other day for 7 days. Scale bar = 50 µm.
E. Quantitative analysis of OPC differentiation by removal of PDGF (n=3 in each group). The stage of differentiation was determined by quantitative analysis of cells immunopositive for myelin basic protein (MBP, red), or neural/glial antigen 2 (NG2, green), expressed as a percent of total cell numbers (DAPI; Blue).
F. Representative images of caspase 3 activation during differentiation of OPCs to GalC+ immature oligodendrocytes. The majority of caspase3 immunopositive cells were also GalC (green) immunopositive (merge is yellow) demonstrating that caspase 3 was activated during cell differentiation in response to TNFα or IL-1β. Scale bar = 50 µm.
G. Quantitation of caspase 3 activation during differentiation of OPCs to GalC+ immature oligodendrocytes. Cell death was quantified as the percent of caspase-3 (red) immunopositive cells (n=6 in each group).

Data information: Data are presented as mean ± S.D. * = p < 0.05, ** = *p* < 0.01, *** = *p* < 0.001 compared to untreated control cultures. ANOVA with Tukey post-hoc comparisons.

### Expression of nSMase2 modifies the cellular response to inflammatory cytokines

Our immunostaining results determined that nSMase2 was expressed in MBP+ oligodendrocytes, but was not expressed in NG2+ OPCs (Fig 3A). Oligodendrocytes exposed to TNFα increased ceramide and decreased sphingomyelin content (Fig 3B), consistent with the known linkage of TNFα to nSMase2 (Adam-Klages, Adam et al., 1996). In contrast, OPCs exposed to TNFα decreased ceramide content and increased sphingosine 1-P (Fig 3B), consistent with a protective response (Brait, Tarrason et al., 2016). These data suggest that the response of OPCs and oligodendrocytes to inflammatory cytokine challenge may be regulated by the developmental expression of nSMase2. When we expressed nSMase2 in NG2+ OPCs these cells became sensitive to TNFα-induced apoptosis, and this cell death was blocked by simultaneous addition of the nSMase2 inhibitor Altenusin (Fig 3D-F); expression of a control EGFP vector in NG2+ OPCs did not modify sensitivity to TNFα-induced apoptosis (Fig 3F). We then reduced expression of nSMase2 in mature oligodendrocytes by lentiviral delivery of a sh-nSMase2-GFP directed against nSMase2 (or a scrambled sh-RNA-GFP), and found that oligodendrocytes were protected from TNFα-induced apoptosis (Fig 3G-I). These data suggest that the developmentally regulated expression of nSMase2 modifies the cellular response to TNFα.

**Fig. 3.**
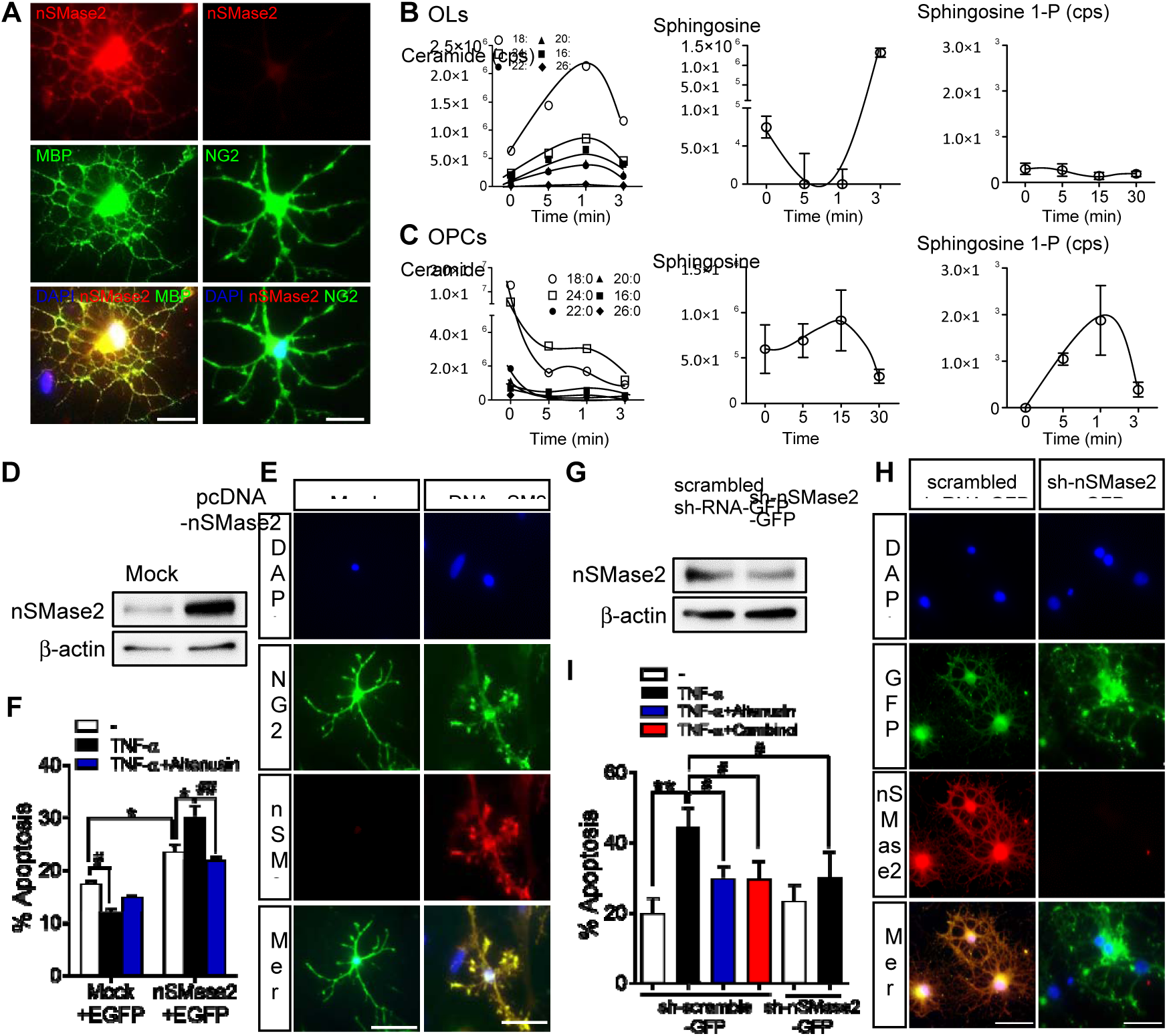
Manipulation of nSMase2 expression regulates cell survival in response to TNFα. A. Representative images showing the expression of nSMase2 (red) in mature oligodendrocytes (OLs; MBP, green), and a lack of nSMase2 expression in oligodendrocyte progenitor cells (OPCs; NG2, green) (co-localization appears as yellow). Scale bar = 20 µm.
B. Quantitative analysis of sphingolipids in OLs following treatment of TNFα (100 ng/ml for the indicated time points) showing increases of ceramide, and decreases in sphingosine (n=3 in each time point).
C. Quantitative analysis of sphingolipids in OPCs following treatment of TNFα (100 ng/ml for the indicated time points) showing decreases ceramide, and increases sphingosine 1-phosphate (n=3 in each time point).
D. Representative immunoblots of overexpression of nSMase2 in OPCs transfected with empty vector (Mock) or pcDNA-nSMase2.
E. Representative fluorescent images of NG2+ OPCs (green) transfected with an empty vector (Mock), or a vector expressing nSMase2 (pcDNA-nSMase2; red). Nuclei were stained with DAPI (blue). Scale bar = 50 µm.
F. Quantitative analysis of apoptosis in OPCs transfected with empty vector or pcDNA-nSMase2 (co-transfected with EGFP-C2 as transfection indicator) followed by treatment of TNFα (100 ng/ml) (n=3 in each group). Apoptotic nuclei (fragmented or condensed) were quantified and expressed as % of EGFP immunopositive cells. Inhibition of nSMase2 with altenusin (25 µM) confirmed that nSMase2 expression in OPCs regulated TNFα induced apoptosis of pcDNA-nSMase2 cells. G Representative immunoblot of knockdown of nSMase2 in OLs transduced with a control lentivirus expressing scrambled RNA (scrambled sh-RNA-GFP), or shRNA directed against nSMase2 (sh-nSM2-GFP).
G. Representative fluorescent images of OLs transduced with a control lentivirus expressing scrambled RNA (scrambled sh-RNA-GFP), or shRNA directed against nSMase2 (sh-nSM2-GFP). Scale bar = 50 µm.
H. Quantitation of apoptotic (fragmented or condensed) nuclei in OLs transduced with the indicated vectors and treated with vehicle, TNFα (100 ng/ml), TNFα + Altenusin (25 µM), or TNFα + cambinol (10 µM) for 24 h (n=3 in each group). Apoptotic nuclei were expressed as % of GFP+ cells.

Data information: Data are presented as mean ± S.D. * = p < 0.05, ** = *p* < 0.01, and # = p < 0.05 as indicated. ANOVA with Tukey post-hoc comparisons.

### Inhibition of nSMase2 protects from cuprizone-induced demyelination

We next determined if inhibition of nSMase2 protected the integrity of white matter during demyelination. Twenty-six days of a cuprizone containing diet led to a 41 ± 9.2% increase in nSMase2 activity in the pCC of mice, an effect that was blocked by unilateral intraventricular infusion of cambinol (a potent inhibitor of nSMase2(Figuera-Losada, Stathis et al., 2015)) 2 days prior to initiating the cuprizone containing diet. Intraventricular infusion of cambinol for 28 days did not change health status of the mice compared to vehicle. Elevations of multiple ceramides (C16:0, 20:0, 22:0, 24:0, 24:1; Monohex-Cer C18:0) in pCC were likewise blocked by ventricular infusion of cambinol in cuprizone-fed mice (Fig 4C). However, decreases in multiple sphingomyelins and sulfatides in cuprizone-fed mice were not modulated by cambinol infusion (Fig 4C). Black gold staining of myelin in the pCC showed a partial protection by cambinol (59 ± 18.3% reduction in staining compared to control) from the reduction in staining in cuprizone-induced demyelination (89 ± 9.5% compared to control, Fig 4D, E).

**Fig. 4.**
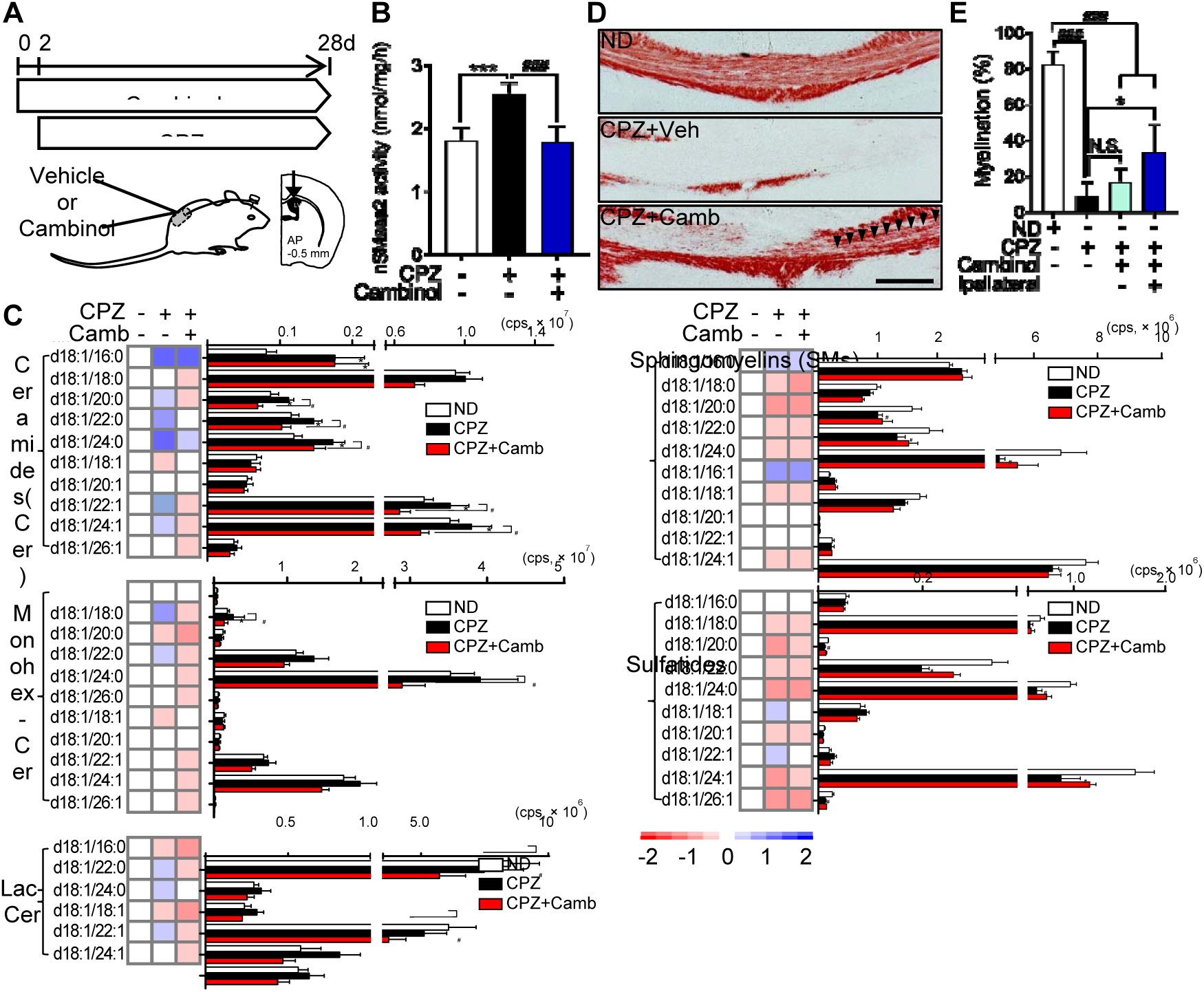
Inhibition of nSMase2 blocked cuprizone-induced demyelination. A. Schematic illustration showing the timing of cambinol infusion during cuprizone (CPZ)-induced demyelination. Cambinol was unilaterally infused into lateral ventricle 2 days prior to cuprizone feeding (CPZ) for 28 days at the rate of 0.31 µg/kg/day.
B. Quantitation of nSMase2 activity in the cortex of mice exposed to CPZ following vehicle or cambinol infusion showing that cambinol blocked CPZ-induced nSMase2 activity (n=5 in each group).
C. Heat maps and quantitative analysis of the indicated class and species of sphingolipids in the pCC of mice fed a standard diet (Normal, n=6)), mice fed CPZ with vehicle infusion (n=6), and mice fed CPZ+cambinol (n=5). CPZ-induced Increases of ceramide were blocked by cambinol infusion.
D. Representative images of black gold stained myelin in the pCC of mice from the indicated treatment groups. Arrowheads show the region of cambinol-mediated protection against CPZ-induced demyelination. Scale bar = 200 µm.
E. Quantification of black gold stained area in the pCC showing a partial protection from demyelination in the CPZ+cambinol group (n=5 in each group).

Data information: Data are presented as mean ± S.D. * = *p* < 0.05, ** = *p* < 0.01, *** = p < 0.001, and # = p < 0.05, ### = p < 0.001 as indicated. ANOVA with Tukey post-hoc comparisons.

### Normalization of myelin ceramide content by inhibition of nSMase2

Modifications in myelin lipid content were readily apparent up to 6 weeks following the return of mice to a normal diet (i.e. during remyelination; see Fig 1). We next determined if inhibition of nSMase2 during the remyelination process modified the lipid content of myelin. Mice were fed a cuprizone containing diet for 28 days followed by unilateral intraventricular infusion of cambinol for 28 days after return to a normal diet (Fig 5A). Activity of nSMase2 was elevated after feeding a diet containing cuprizone for 28 days, and remained elevated for an additional 28 days during remyelination (Fig 5B). Infusion of cambinol during remyelination normalized nSMase2 activity (Fig 5B), and pCC ceramide content, but did not modify increases of sphingomyelin and reductions of sulfatides (Fig 5C). Similar patterns were apparent in multiple other brain regions, albeit to a lesser extent than pCC (Supplementary Fig 2). Although, gross remyelination of pCC cambinol infusion, as determined by black gold staining (Fig 5D,E), did not change significantly, the treatment increased the number of myelinated fibers by 31 ± 10.8% (Fig 5F,G), and the thickness of myelin as evidenced by a 5.0 ± 3.2% reduction in the g-ratio (Fig 5H).

**Fig. 5.**
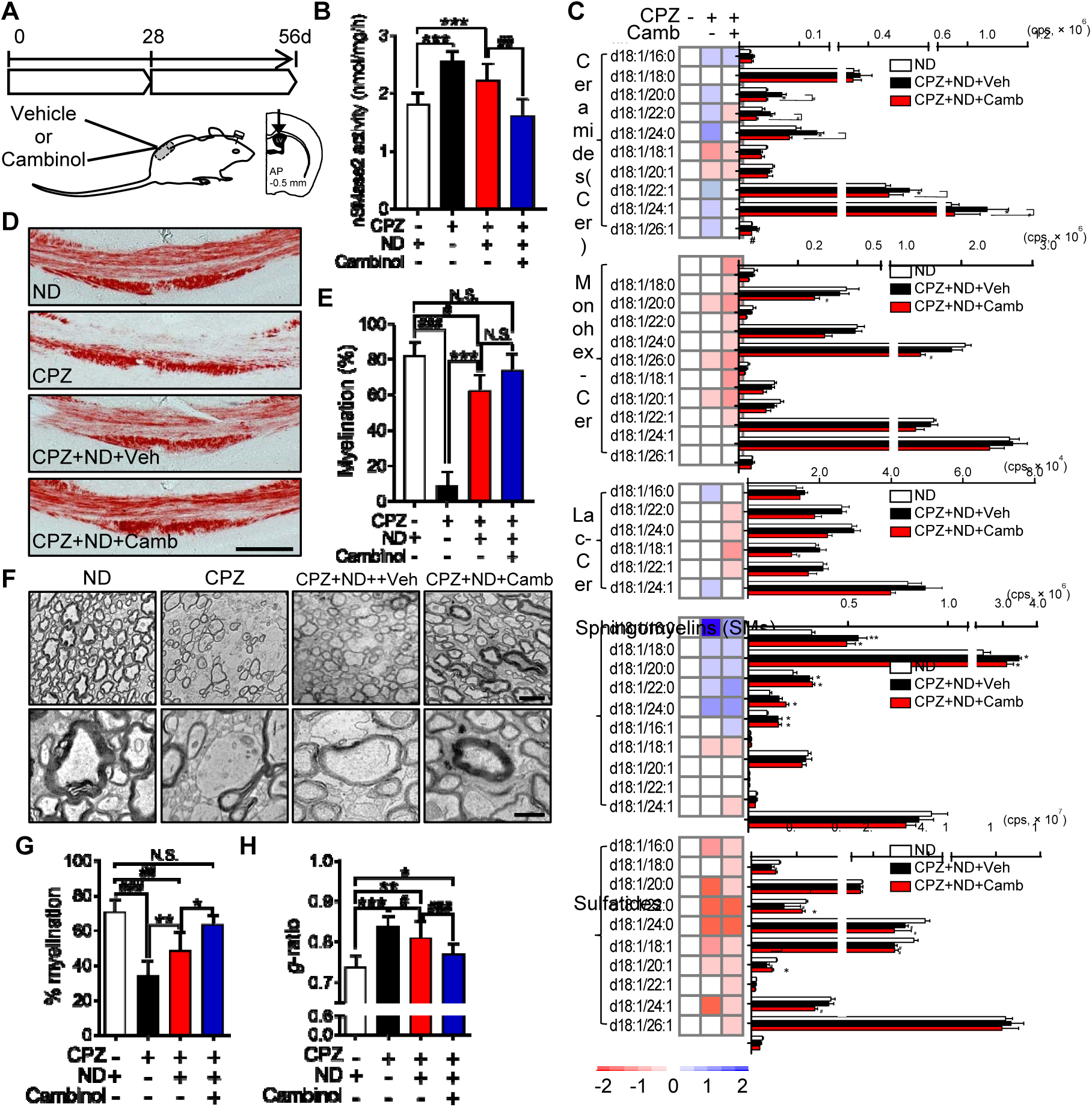
Pharmacological inhibition of nSMase2 enhanced remyelination and improved myelin compaction. A. Schematic illustration showing the timing of cambinol treatment following cuprizone-induced demyelination. After 4-weeks of a cuprizone (CPZ) containing diet, cambinol was unilaterally infused at the rate of 0.31 µg/kg/day for 28 days into the lateral ventricle during return to a normal diet (ND).
B. Quantitation of nSMase2 activity in the cortex of mice fed a ND, a CPZ containing diet for 4-weeks, after return to a ND for 4-weeks (Remyel), and 4-weeks after return to a ND with cambinol infusion showing that cambinol infusion blocks CPZ-induced upregulation of nSMase2 activity (n=5 in each group).
C. Heat maps and quantitative analysis of the indicated class and species of sphingolipids in pCC of mice from the indicated treatment groups (n=5 in each group). Cambinol infusion blocks CPZ-induced upregulation of multiple ceramides, but does not modify increases of sphingomyelins, or decreases in the sulfatide content of pCC.
D. Representative images of black gold staining in the pCC of mice following the indicated treatments. Scale bar = 200 µm.
E. Quantitative analysis of black gold staining in pCC of mice following the indicated treatments (n=5 in each group).
F. Representative electron microscopy images of axons in the pCC of mice following the indicated treatment conditions. Scale bars = 2 μm top panel and 500 nm bottom panel
G. Quantitation of myelinated axons in pCC by electron microscopy showing a larger percent of myelinated axons in CPZ+ND+Camb mice compared to CPZ+ND+Veh (n=100-250 axons from 3 animals per group).
H. Analysis of g-ratios in pCC showing thicker compact myelin structures in CPZ+ND+Camb mice compared to CPZ+ND+Veh (n=100-250 axons from 3 animals per group).

Data information: Data are presented as mean ± S.D. * = *p* < 0.05, ** = *p* < 0.01, *** = *p* < 0.001, and # = p < 0.05, ## = p < 0.01, and ### = p < 0.001 as indicated. ANOVA with Tukey post-hoc comparisons.

### Genetic deletion of nSMase2 in remyelinating oligodendrocytes stabilized sphingolipid composition and promoted remyelination

To conditionally delete smpd3 specifically in a cell-type specific manner, floxed smpd3 mice (*smpd3^fl/fl^*) were crossed with *PDGFRα-CreER* mice (Fig 6A) to selectively delete the nSMase2 gene in remyelinating oligodendrocytes. *PDGFRα-CreER; smpd3^fl/fl^* mice were fed a cuprizone containing diet for 4 weeks, then were administered tamoxifen (1 mg, I.P.) during the first 5 days of return to a normal diet (4 weeks). Mice were sacrificed 4 weeks after return to a regular diet (Fig 6B). Tamoxifen-induced deletion of nSMase2 in remyelinating axons in the *PDGFRα-CreER; smpd3^fl/fl^* mice was validated by immunohistochemical analysis of pCC (Fig 6C). *PDGFRα-CreER; smpd3^fl/fl^* mice fed a cuprizone containing diet showed extensive loss of MBP+ myelin in the pCC compared to the mice fed a standard diet. Additionally, expression of nSMase2 was diminished along with loss of MBP in pCC, indicating that nSMase2 was enriched in MBP+ oligodendrocytes. Following return to a standard diet MBP+ myelin largely recovered in pCC, and was enriched with nSMase2+ cells. Mice in which nSMase2 was deleted by administration of tamoxifen during remyelination (CPZ-ND-Tam) exhibited complete loss of nSMase2 and almost complete recovery of MBP+ myelin in the pCC, indicating that inhibition of nSMase2 promoted remyelination. We next analyzed sphingolipid contents in these mice to confirm if deletion of nSMase2 stabilized sphingolipid perturbations during remyelination (Figure 6C). Similar to the results from wild-type mice exposed to CPZ followed by remyelination, *PDGFRα-CreER; smpd3^fl/fl^* mice subjected to remyelination following CPZ exhibited elevation of multiple ceramides in pCC compared to control mice on a normal diet (C16:0, 20:0, 22:0, 24:0). However, mice administered tamoxifen during remyelination (CPZ-ND-Tam) had lower elevations of these multiple ceramides in pCC compared to Tam- mice (C18:0, 20:0, 22:0, 24:0, 22:1, 24:1). Several monohexosyl ceramides (C24:0, 22:1, 24:1) and lactosyl ceramides (C22:0, C24:1) in pCC of mice subjected to remyelination (CPZ+ND) were different from levels in pCC of control mice, and deletion of nSMase2 significantly lowered their levels during remyelination. Overall levels of sphingomyelins were not significantly altered in mice where nSMase2 was deleted during remyelination. Moreover, the sustained overall decrease of multiple sulfatides in pCC of the mice observed during remyelination was not affected by deletion of nSMase2 during remyelination. Only one species of sulfatide (C18:0) was recovered by deletion of nSMase2 during remyelination. Genomic deletion of nSMase2 in remyelinating oligodendrocytes resulted in an overall decrease of most of ceramides in other brain regions of *PDGFRα-CreER; smpd3^fl/fl^* mice compared to the mice with functional nSMase2 (Supplementary Figure 3). These data suggest that genomic deletion of nSMase2 from remyelinating oligodendrocytes stabilized sphingolipid metabolism during remyelination.

**Fig. 6.**
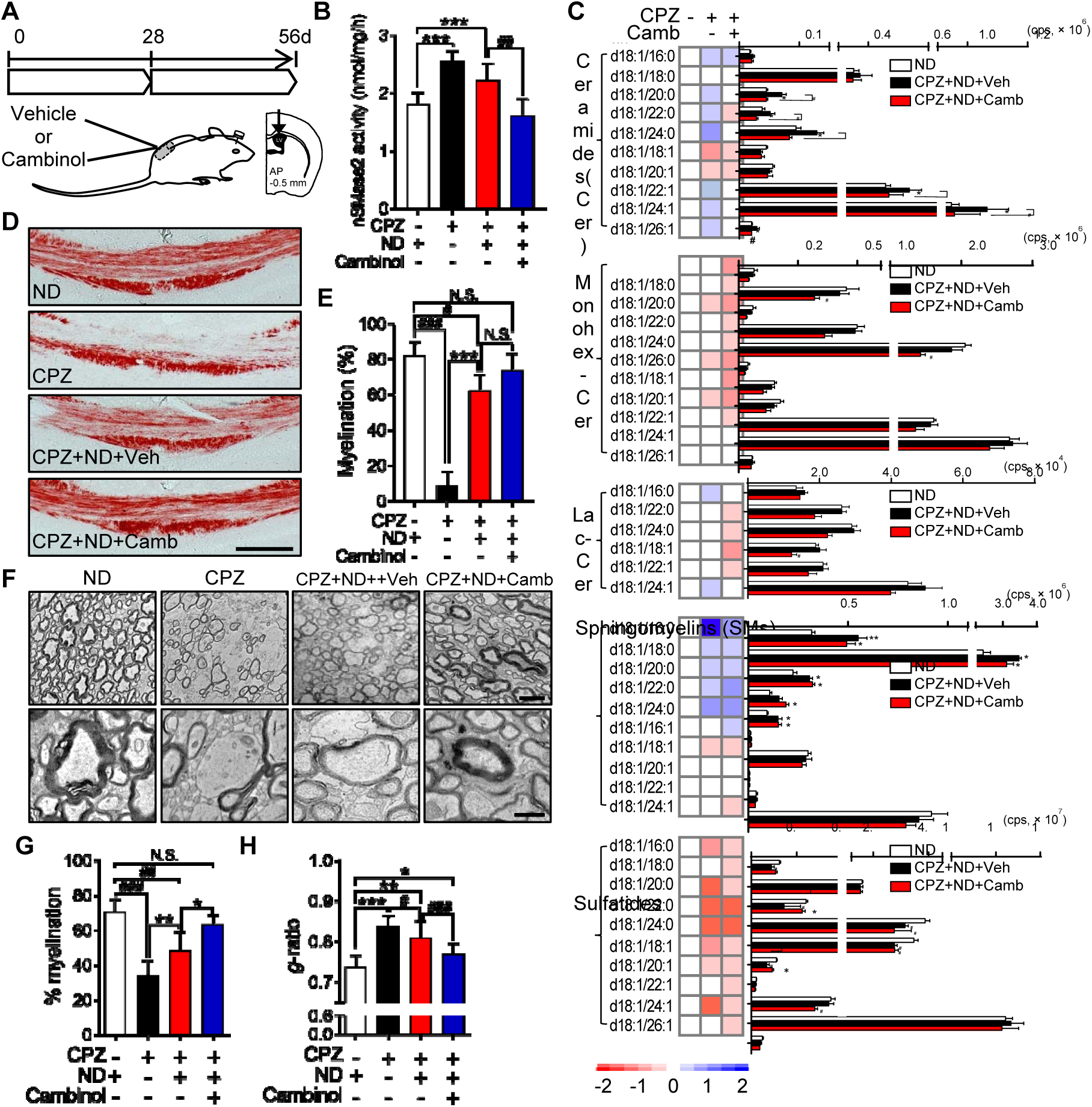
Genetic deletion of nSMase2 during remyelination improves myelination and myelin compaction. A. Schematic illustration for creating PDGFRα-CreER-smpd fl/fl mice.
B. Schematic illustration showing the timing of cuprizone feeding, and tamoxifen treatment (1mg/day, over 5-days) during the return to a normal diet (ND).
C. Representative images of MBP (green) and nSMase2 (red) fluorescence in the pCC of mice fed a normal diet (ND), a cuprizone containing diet (CPZ) for 4-weeks, a CPZ diet and return to ND for 4 weeks (CPZ+Re), and CPZ with tamoxifen induction of nSMase2 deletion during the first 5 days of return to ND (CPZ+ND+Tam). Increased expression of nSMase2 in the CPZ+ND group is absent in the CPZ+ND+Tam group, and myelin basic protein (MBP) staining is more robust in the CPZ+ND+Tam group compared to the CPZ+ND group. Scale bar = 100 μM.
D. Heat maps and quantitative analysis of the indicated class and species of sphingolipids in pCC of mice from the indicated treatment groups (n=4-5 per group). Ceramide content of the pCC was reduced, but sphingomyelins remained elevated and sulfatides reduced in the CPZ+ND+Tam mice compared with the CPZ+ND mice.
E. Representative electron microscopy images of pCC in mice treated with indicated conditions. Scale bars = 2 μm top panel and 500 nm bottom panel.
F. Quantitation of myelinated fibers in pCC showing a greater percent of myelinated fibers in the CPA+ND+Tam mice compared with the CPZ+ND mice (n=100-250 axons from 3 animals per group).
G. Analysis of g-ratio’s for the pCC of mice showing thicker and compact myelin structures in the CPZ+ND+Tam mice compared with the CPZ+ND mice (n=100-250 axons from 3 animals per group).

Data information: Data are presented as mean ± S.D. * = p < 0.05, ** = p < 0.01, *** = p < 0.001, and # = p < 0.05, ## = p < 0.01 as indicated. ANOVA with Tukey post-hoc comparisons.

Next, we analyzed ultrastructure of myelin around axons from mice with deleted nSMase2. While the axons in pCC of *PDGFRα-CreER; smpd3^fl/fl^* mice subjected to remyelination following demyelination (CPZ-ND) were less-myelinated with thinner myelin than mice on normal diet, the axons in nSMase2 deleted mice (CPZ-ND+Tam) were almost completely remyelinated (28.0 ± 10.6% increase over Tam-) and sheaths were similar in thickness to control mice (6.6 ± 5.8% increase than Tam-; Figure 6E-G). These results suggest that nSMase2 expression by remyelinating oligodendrocytes is detrimental for proper remyelination in CPZ-induced demyelination/ remyelination, and that normalizing sphingolipid perturbations by deletion of nSMase2 can enhance remyelination.

## Discussion

Following a demyelinating event, OPCs are recruited to the lesion site where they differentiate into myelinating oligodendrocytes. Myelination requires that oligodendrocyte processes wrap axons and ultimately compact to stabilize the myelin sheath and underling axon. The process of myelination involves a highly synchronized production of multiple types of lipids that regulate membrane curvature and compaction. Here we provide evidence that this orchestrated synthesis of lipids is perturbed during remyelination through mechanisms that involve the sphingomyelin hydrolase nSMase2. OPCs do not express nSMase2 and exhibit a protective/instructional response to TNFα and IL-1β. As OPCs differentiate into MBP+ myelinating oligodendrocytes they express nSMase2 and exhibit an injury response to TNFα and IL-1β. Inhibition of nSMase2 during remyelination in the cuprizone model of demyelination/remyelination restores the ceramide content of myelin to baseline levels, increases the number of remyelinated axons and the thickness of myelin.

The cuprizone diet rodent model of demyelination and remyelination is characterized by a global degeneration of myelin largely in the corpus callosum, with some involvement of cortical and subcortical regions (Matsushima & Morell, 2001). Although the precise mechanism(s) of oligodendrocyte susceptibility to cuprizone remain unclear, and it is not considered a model involving a fulminant immune activation, there is considerable evidence for an immune component to cuprizone-induced demyelination and remyelination. The chemoattractant responsive C-X-C chemokine receptor type 2 on neutrophils is involved in cuprizone-induced demyelination, and inhibition of CXCR2 results in better myelin repair, presumably by allowing optimal spatiotemporal positioning of OPCs in demyelinating lesions (Liu, Darnall et al., 2010). The activation of astrocytes and microglia is readily apparent in the cuprizone model, and appears to involve the Toll-like receptor 2(Esser, Gopfrich et al., 2018). Glial activation is part of the innate immune response, and is critical for remyelination. Activated astrocytes produce the chemoattractant CXCL10 that recruits microglia to phagocytosis cellular debris during demyelination and remyelination. This clearance activity, in conjunction with the increased production of TNF-α, IGF-1, and FGF-2 creates a microenvironment that supports regeneration (Skripuletz, Hackstette et al., 2013, Voss, Skuljec et al., 2012). Ablation of astrocytes in a GFAP-thymidine kinase transgenic mouse model was associated with a failure to clear myelin debris and deficits in remyelination (Skripuletz et al., 2013). Genetic ablation of CXCR3 (receptor for CXCL10 and CXCL9) likewise resulted in deficits of glial activation, debris clearance and remyelination (Krauthausen, Saxe et al., 2014). Similar results were observed in IL-1β-deficient mice that failed to remyelinate properly, and showed a profound delay of OPCs to differentiate into mature oligodendrocytes (Mason, Suzuki et al., 2001). These data suggest that inhibiting the early neuroinflammatory response is detrimental to remyelination. While the inflammatory response is beneficial and necessary for tissue repair following damage, prolonged inflammation can be detrimental to tissue repair (Reviewed in(Le Thuc, Blondeau et al., 2015)). Our data suggest that the developmental expression of nSMase2 may be one mechanistic explanation for this dual effect of inflammation. OPCs do not express nSMase2, and when exposed to TNFα or IL-1β show reduced levels of active caspase3, increased motility, reductions in ceramide, and increases in S1P, each indicative of a protective response. During differentiation, maturing oligodendrocytes express nSMase2, and this expression dramatically changes the cellular response to inflammatory cytokines. Mature oligodendrocytes exposed to TNFα show increased levels of active caspase3, and increased ceramide, indicating a toxic response to this inflammatory cytokine. TNF and IL1 receptors are physically associated with nSMase2 through the WD-repeat protein Fan (Factor associated with nSMase activation) and may form a complex with other modulators including EED (embryonic ectodermal development), and RACK1 (Receptor of activated protein C kinase 1) (Adam-Klages et al., 1996, Taupin, 2010, Tcherkasowa, Adam-Klages et al., 2002). Expressing nSMase2 in OPCs, and genetic deletion of nSMase2 in mature oligodendrocytes confirmed that this differential response to TNFα was due to the expression (or lack of expression) of nSMase2. Thus, expression of nSMase2 during transition of OPCs to oligodendrocytes provides a promising therapeutic target, which does not modify the early protective response of OPCs to inflammatory cytokines, but could protect maturing oligodendrocytes during remyelination.

Myelin has a very high lipid to protein content with the lipid components representing ∼80% of the total dry weight (O’Brien & Sampson, 1965, Rouser, Galli et al., 1965). Maintaining the correct lipid composition of myelin is essential for myelin structure and function. The lipid content of CSF and normal appearing white matter in MS brain tissues is modified with a higher phospholipid and ceramide content, increased lipid peroxidation, and a total sulfatide content that is reduced by 25% (Takahashi & Suzuki, 2012, Vidaurre, Haines et al., 2014, Wheeler et al., 2008). Remyelinated axons in the cuprizone model recapitulate some of these modifications in lipid content. We found that remyelination was incomplete and many of the remyelinated axons in the posterior corpus callosum were thinly myelinated, or the myelin was disorganized and not compacted, similar to findings by other groups (Guo, Suo et al., 2018, Mullin, Cui et al., 2017, Tagge, O’Connor et al., 2016). This disorganization was accompanied by an abnormal lipid content with increased ceramide, increased sphingomyelin, reduced hexosylceramide and sulfatide. Pharmacological inhibition or a targeted knock-out of nSMase2 in newly generated oligodendrocytes corrected the ceramide content of remyelinated axons in the posterior corpus callosum, increased the myelin thickness, and dramatically reduced the number of axons with disorganized/uncompacted myelin. However, inhibition of nSMase2 did not normalize the reduced level of GalCer or sulfatide content of remyelinated axons.

GalCer, and sulfatide account for ∼20% and 5% respectively of myelin lipid content (Boggs, Gao et al., 2008, Marcus, Honigbaum et al., 2006). The synthesis of these galactosphingolipids involves the addition of galactose from UDP-galactose to ceramide in a reaction catalyzed by ceramide galactosyltransferase, and the subsequent addition of a sulfate group by cerebroside sulfotransferase (CST) (Honke, 2013). The synthesis of GalCer and sulfatide occurs in mature myelinating oligodendrocytes, and marks the transition from late OPCs to immature oligodendrocytes (Poduslo & Miller, 1985, Raff, Mirsky et al., 1978). Both GalCer and sulfatide form phase separated domains in model membranes, and are strongly associated with ceramide and cholesterol in lipid raft domains (Hao, Sun et al., 2009). The importance of these lipids in myelin stability was demonstrated in CGT-/-knock out mice that do not make GalCer or sulfatide. CNS myelin in these mice is thinner, nodal long is increased and lateral loops are widely spaced, with extensive vacuolization between the sheaths and axolemma (Bosio, Binczek et al., 1998, Dupree, Coetzee et al., 1998). CGT-/-mice typically die within three months of age (Dupree et al., 1998). CST-/-knockout mice are unable to synthesize sulfatide, but the level of other glycolipids including GalCer normal. These mice are born healthy and produce compact myelin (albeit thinner myelin), but display progressive myelin abnormalities including a reduction in total myelin lipid content (due to a reduction in lipid synthesis by oligodendrocytes), nodal structure deterioration, myelin vacuolar degeneration, and reductions of axon caliber, with abnormal clustering of Na^2+^ and K^+^ channels(Ishibashi, Dupree et al., 2002, Marcus et al., 2006, Palavicini, Wang et al., 2016). These findings, in consideration of our current results showing reduction of GalCer and sulfatide in remyelinated axons, suggests there may be a deficiency in CST activity during remyelination.

Our findings indicate that developmental expression of nSMase2 modifies the cellular response to inflammation, from being protective at the OPC stage, to damaging in myelinating oligodendrocyte stage. Pharmacological inhibition or genetic deletion of nSMase2 did not modify the instructive or protective response of OPCs to inflammatory stimuli, but protected myelinating oligodendrocytes. This targeted protection of maturing oligodendrocytes partially restored myelin lipid composition, and improved the biophysical properties of myelin to allow greater compaction, suggesting that targeted modification of this pathway within premyelinating oligodedendrocytes could be beneficial in promoting myelin repair in MS.

## Materials and Methods

### Animals and induction of demyelination / remyelination

Pregnant female Sprague-Dawley (SD) rats (embryonic day 17), and male C57BL6 mice (8-10 weeks old) were obtained from the Jackson Laboratory (Bar Harbor, ME). Mice were housed in a temperature and humidity-controlled room under a 12h light cycle. Mice were allowed to acclimate to the colony room for at least 7 days after arrival before experimentation. All procedures were conducted in accordance with NIH guidelines for the Use of Animals and Humans in Neuroscience Research and approved by Institutional Animal Care and Use Committee (Johns Hopkins University School of Medicine). To induce demyelination, mice were fed 0.2% (w/w) cuprizone (bis(cyclo-hexanone) oxaldihydrazone (Sigma) mixed with a powdered rodent diet containing18% protein (Teklad Global) for 4 weeks. Mice were then returned to a normal diet (ND) for 4-6 weeks to promote remeylination (173 mice total; 52 mice for ND, 121 mice for cuprizone diet followed by direct sacrifice, ND, or ND with drug infusion. Mice were identified by earmarks and numbered accordingly, randomly grouped before starting demyelination or remyelination. During experiments and analysis, the investigators were blinded to experimental group.

*In vivo* inhibition of nSMase2 was accomplished by a mini-osmotic pump (Alzet, Cupertino, CA) infusion of cambinol (Figuera-Losada et al., 2015) into the lateral ventricle (0.31 mg/day, 0.11 µl/h flow rate; AP, -.0.5 mm; ML, 1.0 mm; DV, 2.5 mm). Drug delivery began 2 days prior to initiation of the cuprizone diet, or after 4 weeks of the cuprizone diet. At the end of experimental protocols brains were rapidly removed and dissected on ice to isolate posterior corpus callosum (pCC), anterior corpus callosum (aCC), cerebral cortex (CTX), hippocampus, striatum (STR) and cerebellum (CBL). Tissues were flash frozen and stored at −80C.

#### Conditional knock out of nSMase2

The mouse embryonic stem cells (ES) containing a conditional knock-out allele for the smpd3 gene (nSMase2) was obtained from European Conditional Mouse Mutagenesis Program (EUCOMM, ES cell clone, HEPD0746_4_A11; targeting vector, PG00238_Z_4_F05). The ES cells harbored a LacZ reporter cassette in exon 1, resulting in expression of the β-galactosidase enzyme under control of the endogenous smpd3 promoter. The LacZ cassette was flanked by FRT sites that allows deletion of the LacZ and the stop codon when crossed with Flippase expressing (Flp-deletor) mice. Heterozygote male ES cells were injected into albino C57B6 embryos (JHU transgenic core facility), and implanted into females. The resultant male chimeric mice were crossed with albino C57B6 mice. Albino coated offspring were genotyped for the presence of the LacZ gene (forward primer, 5’-CGATCGTAATCACCCGAGTGT-3’; reverse primer, 5’-CCGTGGCCTGACTCATTCC-3’; reporter, 5’-CCAGCGACCAGATGAT-3’, Transnetyx). Male mice containing the LacZ gene were then crossed with a homozygeous *flpo* C57B6N mouse (Mutant Mouse Resource & Research Centers, NIH) to delete the LacZ gene. These mice were back-crossed with a non-genetically modified C57B6N (Charles River, Frederick, MD) strain to remove the FRT gene. Floxed smpd3 was confirmed by genotyping using wild-type smpd3 primers (forward primer, 5’-TCTTTCCTGGTCTAGTTGGCACTA-3’; reverse primer, 5’-GAGCCAGGGATGTGTTTAAAGTG-3’; reporter, 5’-TAGAGGAGCTGCAAACAT-3’) and floxed smpd3 primers (forward primer, 5’-GCTGGCGCCGGAAC-3’; reverse primer, 5’-GCGACTATAGAGATATCAACCACTTTGT-3’; reporter, 5’-AAGCTGGGTCTAGATATC-3’). Female *smpd3^fl/f^* mice were crossed with male hemizygous *PdgfaR-CreER* (Kang, Fukaya et al., 2010)(Jackson, Bar Harbor, Maine) to generate *PdgfaR-CreER*; *smpd3^fl/fl^* mice. The presence of cre recombinase in *PdgfaR-CreER*; *smpd3^fl/fl^* mice was confirmed by genotyping (forward primer, 5’-TTAATCCATATTGGCAGAACGAAAACG-3’; reverse primer, 5’-CAGGCTAAGTGCCTTCTCTACA-3’; reporter, 5’-CCTGCGGTGCTAACC-3’). Two to three months old mice (males and females) were exposed to cuprizone feeding for 4 weeks to induce demyelination and recovered for 4 weeks with normal diet (34 mice total; 9 mice for ND, 25 mice for cuprizone followed by direct sacrifice, ND, or ND+tamoxifen). For the first 5 days of remyelination the mice were administered 4-hydroxytamoxifen (4-HT, 1mg/day/mouse, Sigma) as previously described (Baxi, DeBruin et al., 2015), to delete *smpd3*. The mice were sacrificed after 4 weeks of remyelination, and the brains were rapidly extracted and stored at −80°C for analysis.

### Histology and Electron Microscopy

Mice were transcardially perfused with 0.9% NaCl followed by 4% PFA, and the brains were post-fixed with 4% PFA for overnight at 4°C. Black Gold staining was conducted according to the manufacturer’s instructions (Black Gold II myelin staining kit, EMD Millipore, Billerica, MA). Briefly, 30μm thick brain sections were incubated with 0.3% Black Gold at 60°C for 20min. Sections were then fixed with 1% sodium thiosulfate for 3min, dehydrated using a series of gradated alcohols, cleared in xylene, and cover-slipped with mounting medium (EMD Millipore, Billerica, MA). Myelination of axons in pCC was calculated as the percent area stained with Black Gold in total pCC from 3 sequential sections (180 µm apart).

For electron microscopy (EM) mice were anesthetized with isofluorane and perfused transcardially with 4% PFA, 0.1% glutaraldehyde, in 0.1M sodium phosphate buffer (pH 7.4). Brains were post fixed with 4% PFA, 2%glutaraldehyde, 2.5% sucrose, and 3 mM NaCl in 0.1M sodium cacodylate buffer (pH 7.4). Fixed tissues were dissected, dehydrated in graded ethanol, embedded in EMBed-812 resin (Electron Microscopy Sciences, Hatfield, PA). Thin sections (90nm) were cut with a diamond knife using a Reichert-Jung Ultracut E ultramicrotome and picked up with copper slot (1 × 2 mm) grids. Grids were stained with 2% uranyl acetate and lead citrate then viewed using Zeiss Libra 120 transmission electron microscope with a Veleta camera (Olympus, Muenster, Germany). G-ratios were calculated as the % of axonal myelination by measuring the inner and outer diameter of myelinated axons(Baxi et al., 2015). Ten images selected from the central posterior corpus callosum were quantified per mouse (∼20-50 axons/ image).

### Measurement of nSMase2 activity

Brain tissues were homogenized in ice-cold Tris-HCl buffer (0.1 M, pH 7.5), containing 250 mM sucrose, 10 mM EGTA (Research Products International, Prospect, IL), 100 µM sodium molybdate, and protease inhibitors. Measurement of nSMase2 activity was determined based on the direct hydrolysis of [^14^C]-SM to [^14^C]-phosphorylcholine and ceramide as previously described(Figuera-Losada et al., 2015). Bovine [N-methyl-14C]-sphingomyelin [^14^C]-SM (10µM, 52 mCi/mmol, Waltham, MA) was evaporated to dryness under a stream of nitrogen for 1h at 45°C. The dried [^14^C]-SM was re-constituted in Tris-HCl buffer (0.12M, pH7.5, Quality Biological Inc Gaithersburg, MD) with 20mM MgCl_2_ (Sigma), 0.02% Triton X-100 (Sigma), and a mixed micelle solution of the substrate that was prepared using a 10sec vortex, a 1min bath sonication, and a 1 min bath incubation at 37°C. The assay was conducted in 8-well PCR tubes strips, arranged in 96-well PCR tube racks. The reaction mixture, containing 25μl of prepared substrate ([^14^C]-SM 10 µM) and 20μl of tissue lysate or recombinant human nSMase2. At the end of the incubation period, the reaction was terminated by the addition of 30μl water and 175μl of chloroform/methanol (2:1, v:v), followed by vigorous vortexing. To achieve phase separation PCR tubes were centrifuged at 2000g for 1h using a Beckman GS-6R centrifuge equipped with a PTS-2000 rotor. A 50 μL aliquot of the upper aqueous phase, containing the liberated [^14^C]-phosphorylcholine, was analyzed for radioactivity using Perkin Elmer’s TopCount instrument in conjunction with their 96-well LumaPlates, normalized to protein content, and data presented as pmol/mg/h.

### Rat primary oligodendrocyte cultures

Rat oligodendrocyte precursor cells (OPCs) were prepared from 3-5 days old Sprague-Dawley rat cortices as described previously (Chen, Balasubramaniyan et al., 2007). Dissociated cells were plated on culture dishes coated with poly-D-lysine (10 µg/ml, Sigma, St Louis, MO) in DMEM containing 15% fetal bovine serum (FBS), 2mM glutamate (Life Technologies, Grand Island, NY), 100μM nonessential amino acids (NEAA, Life Technologies), 1mM sodium pyruvate (Life Technologies), 100U/ml penicillin, 100μg/ml streptomycin, and 10ng/ml platelet-derived growth factor (PDGF, R/D systems, Minneapolis, MN). After 7-10 days in culture, OPCs were collected by shaking (200 rpm overnight) and re-plated in DMEM containing 0.5% FBS, 1% N2 supplement (Life Technologies), 2mM glutamate, 100μM Non-essential amino acid (NEAA), 1mM sodium pyruvate, 100U/ml penicillin, 100μg/ml streptomycin, 0.5mM N-acetylcysteine (Sigma), and 10ng/ml PDGF.

OPCs were differentiated into mature oligodendrocytes (OLs) by removal of PDGF from culture media for 6 days. Cells were fixed with 4% PFA, blocked with 10% normal goat serum (v:v) in PBS containing 0.1% (v:v) Triton X-100 (PBS-T) for 1h at room temperature, then incubated with antibodies against NG2 (EMD Millipore, 1:1,000), MBP (EMD Millipore, 1:1,000) or GalC (EMD Millipore, 1:1,000) overnight at 4°C. Cells were then washed 3 times with PBS-T, and incubated for 1h at room temperature with an Alexa 488 or 594 conjugated secondary antibodies (Thermo Fisher Scientific, Waltham, MA). Approximately 60% of cells were MBP+ after 6 days of differentiation.

### Cell Motility

Primary OPCs cultures maintained in 37°C and 5% CO_2_ in a live imaging microscope and treated with 25 or 100ng/ml of TNF. Their dynamics were monitored continuously by collecting images every 30min for 12h. Images were processed by Image J (NIH), and data expressed as arbitrary units (AU).

### Manipulation of neutral sphingomyelinase 2 expression

#### nSMase2 expression

A pcDNA-nSMase2 vector (4 pg/cell of DNA) was delivered to OPCs by electroporation (model ECM830, Harvard Apparatus, Holliston, MA; 2 mm cuvette, voltage 200 V, pulse length 5 ms, single pulse/ uni-direction). An empty vector (Mock, pcDNA 3.1, Thermo Fisher Scientific) was used as a control. A vector expressing EGFP-C2 (Clontech, Mountain View, CA) was co-electroporated into cells a transfection indicator. Forty-eight hours after electroporation expression of nSMase2 was quantified by immunoblotting, and immunofluorescent staining. Cells were harvested and proteins resolved by 10% SDS-PAGE and transferred to PVDF membrane (BioRad, Hercules, CA). Non-specific binding sites were blocked with 5% (w/v) milk in TBS containing 0.1% Tween 20 (TBS-T), and incubated with an antibody against nSMase2 (1:1,000, Santacruz Biotechnology, Dallas, TX) overnight at 4°C. Following 3 washes with TBS-T the immunoblot was incubated for 1h with an rabbit IgG HRP-linked antibody (1:1,000; Cell signaling technology, Danvers, MA) in TBS-T, and developed by enhanced chemiluminescence. Image analysis was conducted using a G:BOX Imaging system (Syngene, Frederick, MD). For immunostaining, the cells were fixed with 4% PFA, blocked with 10% normal goat serum for 1h at room temperature, and then incubated with an antibody against nSMase2 (1:200) overnight at 4°C. Cells were washed 3 times with PBS-T and incubated for 1 h at room temperature with Alexa 594 conjugated with anti-rabbit IgG secondary antibody.

#### nSMase2 knock-down

Oligodendrocytes were pretreated with polybrene (5 µg/ml) for 10min then infected with lentivirus expressing sh-nSMase2-GFP (sh-nSMase2-GFP, Applied Bio Materials, BC, Canada), or a scrambled RNA lentiviral vector (sc-con-GFP) (both MOI 5.0), and knockdown of nSMase2 was validated by immunofluorescence and immunoblotting.

### Cell Proliferation and Survival

Cell proliferation was quantified in OPCs treated with 0.5mg/ml Bromodeoxyuridine (BrdU, Sigma, St Luis, MO) prior to exposure to TNFα (0-100ng/ml), IL-1β (0-100ng/ml), or vehicle for 2 days. Cells were fixed with 4% PFA, treated with 2N HCl for 30 min, and pH neutralized with sodium borate (pH 9.0) for 10min. Non-specific binding was blocked with 10% normal goat serum in PBS-T, and cells were incubated with an antibody directed against BrdU (1:200, Sigma, St Luis, MO) for 2h at room temperature in PBS. Cells were washed 3 times with PBS, and incubated for 2h at room temperature with an Alexa 594 conjugated secondary antibody in PBS-T (Thermo Fisher Scientific, Waltham, MA). Cells were washed and mounted on glass slides (Vector laboratories, Burlingame, CA).

Cell survival was quantified in OPCs and OLs sixteen hour after treatment with TNF-α (100 ng/ml), or IL-1β (100ng/ml). Nuclei were stained with Hoechst (1μg/ml, Thermo Fisher Scientific), fixed and the number of pyknotic nuclei (apoptotic) expressed as a ratio to the number of diffusely stained nuclei (healthy). In some experiments survival was also determined by immunostaining for active caspase3. Non-specific binding was blocked with normal goat and horse serum in PBS-T for 2h at room temperature. Cells were then washed 3 times with PBS and incubated with an anti-active Caspase-3 antibody (EMD Millipore, Billerica, MA, 1:200) for 2h at room temperature. Proliferation and survival were quantified in a minimum of 150 cells from at-least three-independent experiments by an experimenter blinded to the experimental condition.

### Pharmacological treatments

Cells were pretreated with the nSMase2 inhibitor altenusin (25µM, Enzo Life Sciences Inc, Farmingdale, NY), cambinol (10µM, Sigma) for 30 min prior to experimental treatments.

### Mass spectrometry

A crude lipid extraction of posterior corpus callosum (pCC), anterior corpus callosum (aCC), cerebral cortex (CTX), striatum (STR), hippocampus (HIP), and cerebellum (CBL) was performed using a modified Bligh/Dyer(Bligh & Dyer, 1959) procedure with ceramide and sphingomyelin (SM) d18:0/12:0 included as internal standards (Avanti Polar Lipids, Alabaster, AL, USA)(Haughey, Cutler et al., 2004). The organic layers containing crude lipid extracts were dried in a nitrogen evaporator (Organomation, Berlin, MA, USA) and suspended in MeOH prior to analysis. Chromatographic separations were conducted using a Shimadzu ultra-fast liquid chromatography system (Shimadzu, Nakagyo-ku, Kyoto, Japan) coupled to a C18 reverse-phase column (Phenomenex, Torrance, CA, USA). Eluted samples were injected into an API3000 triple quadrupole mass spectrometer (AB/Sciex, Thornhill, ON, Canada) where individual ceramide and sphingomyelin species were detected by multiple reaction monitoring. Instrument and HPLC parameters have been previously described(Mielke, Bandaru et al., 2015a, Mielke, Bandaru et al., 2015b). Eight-point calibration curves (0.1–1000 ng/mL) were constructed by plotting area under the curve (AUC) for each ceramide calibration standard d18:1/C16:0, d18:1/C18:0, d18:1/C20:0, d18:1/C22:0, d18:1/C24:0, and SM calibrations standards C16:0, C18:0, C20:0, C22:0 and C24:0 (Avanti polar lipids, Alabaster, AL, USA). Correlation coefficients for standard curves were > 0.999. Ceramide and SM concentrations were calculated by fitting the identified ceramide and SM species to these standard curves based on acyl chain length. Intra-day coefficients of variation for each ceramide SM species were less than 10%(Mielke et al., 2015b) Instrument control and quantification were performed using Analyst 1.4.2 and MultiQuant software (AB Sciex Inc. Thornhill, Ontario, Canada. (Lakshmni needs to add sulfatides)

### Statistics

Data were analyzed by One-way analysis of variance (One-way ANOVA) followed by Tukey post-hoc comparisons when group differences were significant (Graphpad software, La Jolla, CA, USA). Results are expressed as mean ± S.D. as indicated.

### Data availability

The raw data that support the findings of this study are available from the corresponding author by request.

## Acknowledgements

These studies were supported by National Institutes of Health, MH110246, DA040390, MH096636 MH105280 to NJH, and R37NS041435 to PAC. Support was also provided from The Hilton Foundation to PAC & NJH, and from the Dr. Miriam and Sheldon G. Adelson Medical Research Foundation DEB.

## Author contributions

The study was designed by S.W.Y. and N.J.H. Th majority of *in vivo* and *in vitro* experiments were performed by S.W.Y. Cuprizone feeding in mice was performed by M.D.S. Creation of PDGF-Cre-Smpd3 mice was performed by A.A. Analyses of electron microscopy were performed by S.-W.Y. and S.S.K. Analyses of nSMase2 activity were performed by A.G.T. All of the LC-MS analyses were performed by E.G.B. and M.M. Manuscript revisions and comments included C.R., B.S.S., D.E.B., and P.A.C.. S.W.Y. and N.J.H. wrote manuscript. All authors read and revised the manuscript.

## The paper explained

### PROBLEM

Following and demyelinating event, oligodendrocyte progenitor cells (OPCs) are recruited to the site of damage and differentiate into myelinating oligodendrocytes to remyelinate axons. However, for reasons that are not completely understood, remyelination is often incomplete, with thin and/or disorganized myelin structures. These incompletely remyelinated axons are susceptible to secondary demyelination.

### RESULTS

The cuprizone model of demyelination/remyelination recapitulates the phenotype of remyelinated axons that often exhibit a thin and disorganized myelin structure. Remyelinated axons contain high amounts of ceramide and sphingomyelin with lower amounts of sulfatides compared to normal myelin. Ceramides and sphingomyelins form highly organized and tightly packed structures that are not amenable to the curvature required for myelin to wrap axons and compact. We found that OPCs do not express the sphingomyelin hydrolase neutral sphingomyelinase-2 (nSMase2) and react to the inflammatory cytokines TNFα and IL-1β with increased motility and a protective molecular phenotype that includes a reduced ceramide content. As OPC differentiate, nSMase2 expression is turned on, and the differentiating oligodendrocytes react to inflammatory cytokines with increased ceramide, and decreased survival. Pharmacological inhibition or targeted genetic deletion of nSMase2 in OPCs partially restored the lipid composition of remyelinated axons, and largely restored myelin thickness and compaction.

### IMPACT

Our findings suggest that inhibiting nSMase2 following a demyelinating event could improve the quality of remyelinated fibers by normalizing the ceramide content. A more organized and compacted myelin structure would be more stable and less prone to secondary demyelination.

---

**Supplementary Figure 1.**
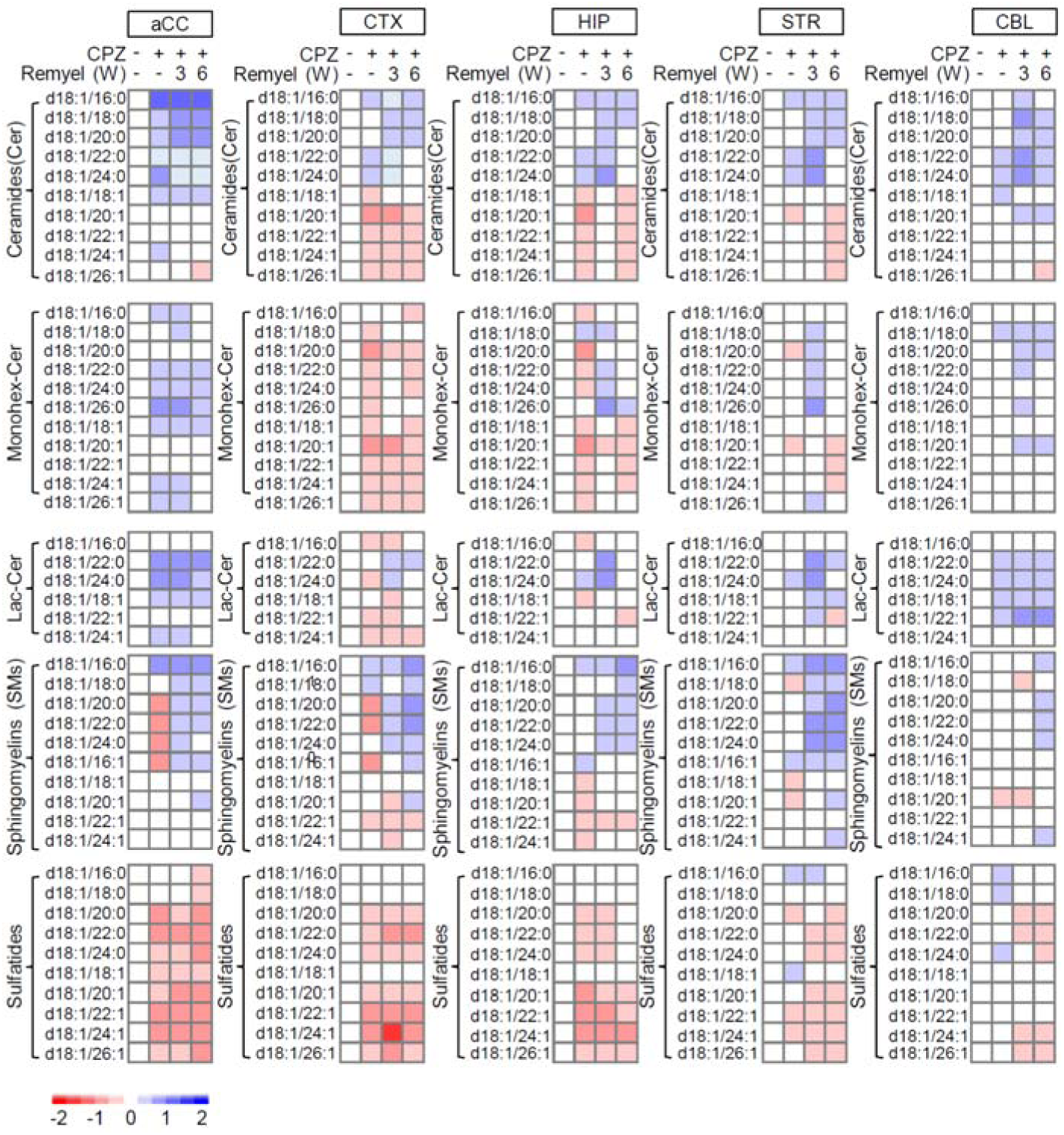
Compositional changes in the sphingolipid content of multiple brain regions after demyelination and during remyelination. Mice were fed either a normal diet, a diet containing cuprizone (CPZ) for 4-weeks, a diet containing CPX followed by return to a normal diet for 3, or 6 weeks. Heat maps showing relative content of the indicated class and species of sphingolipids in posterior corpus callosum (pCC), anterior corpus callosum (aCC), cerebral cortex (CTX), hippocampus (HIP), striatum (STR), and cerebellum (CBL) following the indicated treatment conditions.

**Supplementary Figure 2.**
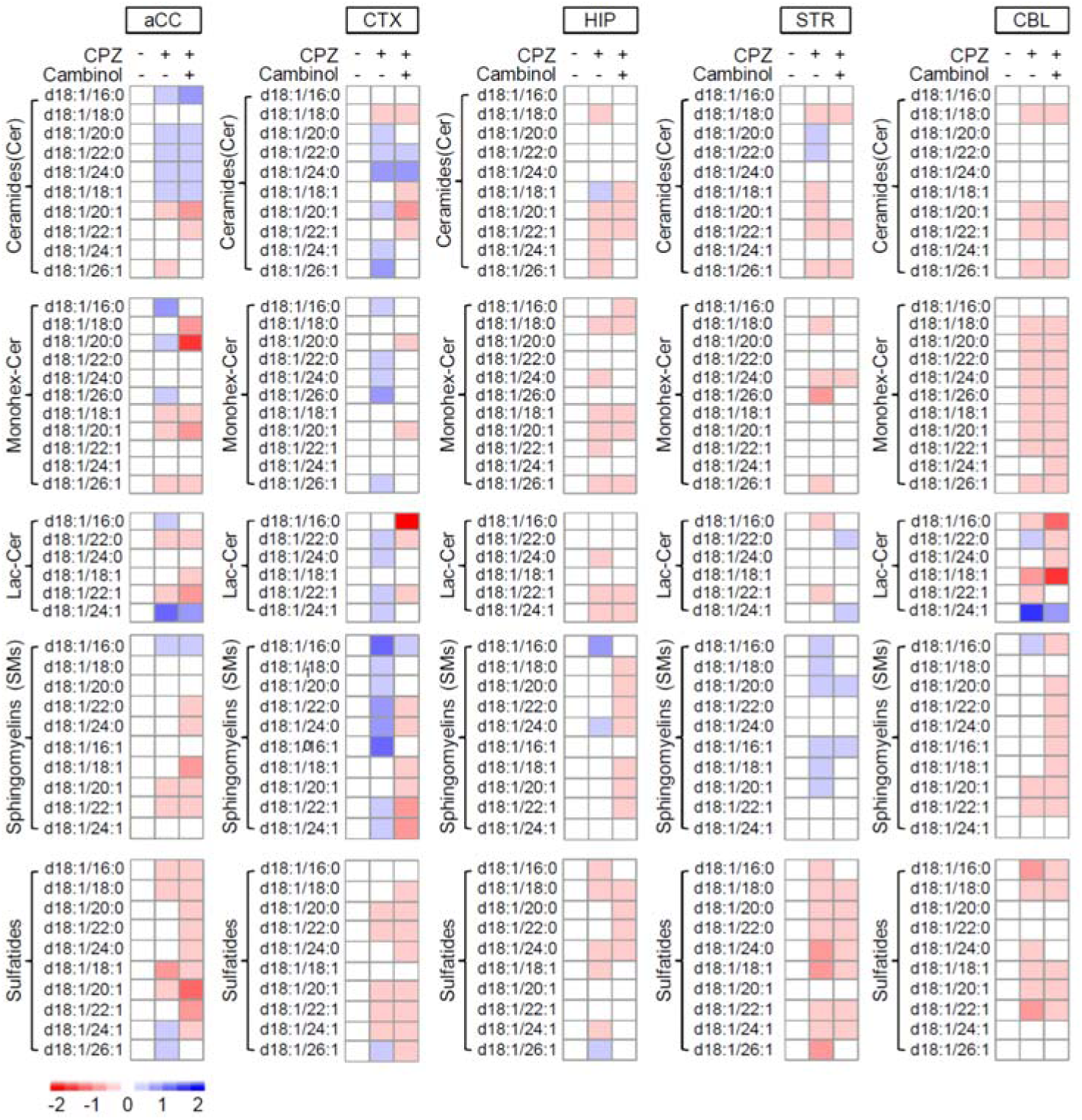
Inhibition of nSMase2 modifies the sphingolipid content of multiple brain regions during remyelination. Heat maps showing the compositional changes in brain sphingolipid content of the posterior corpus callosum (pCC), anterior corpus callosum (aCC), cerebral cortex (CTX), hippocampus (HIP), striatum (STR), and cerebellum (CBL) following cuprizone induced demyelination, remyelination, and remyelination with unilateral infusion of cambinol into the lateral ventricle (28 day infusion at the rate of 0.31 µg/kg/day).

**Supplementary Figure 3.**
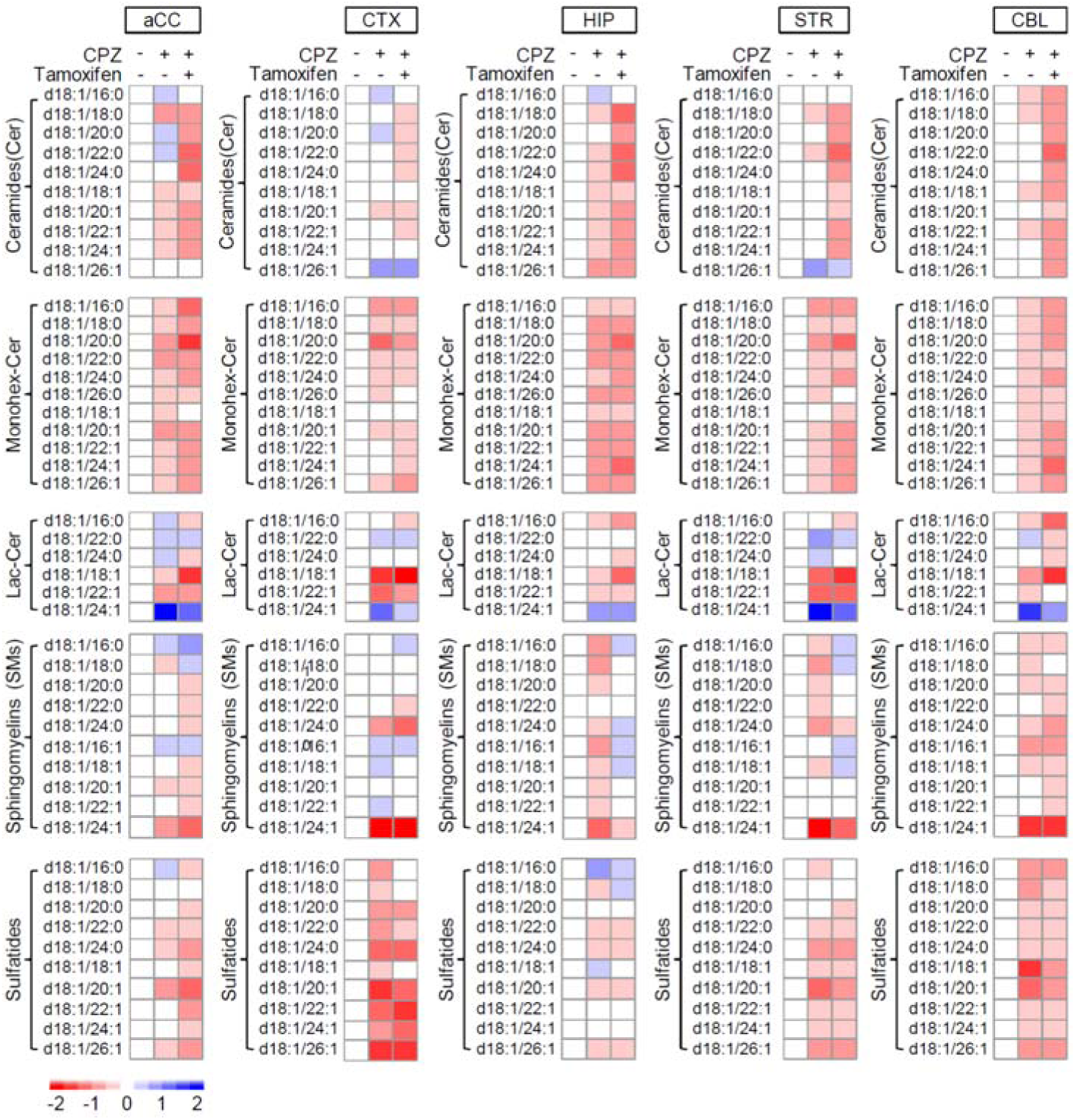
Heat maps of compositional changes in brain sphingolipid content after demyelination and during remyelination in PDGFRα(CreER)-smpd3^fl/fl^ mice. Heat maps showing the compositional changes in brain sphingolipid content of the posterior corpus callosum (pCC), anterior corpus callosum (aCC), cerebral cortex (CTX), hippocampus (HIP), striatum (STR), and cerebellum (CBL) following cuprizone induced demyelination, remyelination, and remyelination following selective deletion of nSMase2 in PDGFRα/NG2 cells.

